# Plasticity impairment alters community structure but permits successful pattern separation in a hippocampal network model

**DOI:** 10.1101/2020.11.30.403766

**Authors:** Samantha N. Schumm, David Gabrieli, David F. Meaney

## Abstract

Patients who suffer from traumatic brain injury (TBI) often complain of learning and memory problems. Their symptoms are principally mediated by the hippocampus and the ability to adapt to stimulus, also known as neural plasticity. Therefore, one plausible injury mechanism is plasticity impairment, which currently lacks comprehensive investigation across TBI research. For these studies, we used a computational network model of the hippocampus that includes the dentate gyrus, CA3, and CA1 with neuron-scale resolution. We simulated mild injury through weakened spike-timing-dependent plasticity (STDP), which modulates synaptic weights according to causal spike timing. In preliminary work, we found functional deficits consisting of decreased firing rate and broadband power in areas CA3 and CA1 after STDP impairment. To address structural changes with these studies, we applied modularity analysis to evaluate how STDP impairment modifies community structure in the hippocampal network. We also studied the emergent function of network-based learning and found that impaired networks could acquire conditioned responses after training, but the magnitude of the response was significantly lower. Furthermore, we examined pattern separation, a prerequisite of learning, by entraining two overlapping patterns. Contrary to our initial hypothesis, impaired networks did not exhibit deficits in pattern separation with either population- or rate-based coding. Collectively, these results demonstrate how a mechanism of injury that operates at the synapse regulates circuit function.

**Author summary:** Traumatic brain injury causes diverse symptoms, and memory problems are common among patients. These deficits are associated with the hippocampus, a brain region involved in learning and memory. Neural plasticity supports learning and memory by enabling the circuit to adapt to external stimulus. After brain injury, plasticity can be impaired, perhaps contributing to memory deficits. Yet, this mechanism of injury remains poorly understood. We implemented plasticity impairment and learning in a network model of the hippocampus that is unique because it has a high degree of biological detail in its structure and dynamics compared to other similar computational models. First, we examined the relationship between neurons in the network and characterized how the structure changed with injury. Then we trained the network with two input patterns to test the function of pattern separation, which is the ability to distinguish similar contexts and underpins general learning. We found that the strength of the encoded response decreased after impairment, but the circuit could still distinguish the two input patterns. This work provides insight into which specific aspects of memory become dysfunctional after injury.

## Introduction

Traumatic brain injury (TBI) is a debilitating condition that involves dysfunction across diverse neural circuitry. Often a result of impacts to the head, TBI is pervasive with up to 2.5 million cases recorded in 2014 [1,2]. The incidence of TBI has risen along with increasing societal awareness of the issue [3], owing in part to the effects of concussion on adolescents and young adults [4]. Despite efforts to mitigate sports-related and other impacts, young people remain affected and can suffer long-term problems from even mild injuries [4–6]. Although standards of diagnosis are improving, treatments for TBI are lacking [7]. Fortunately, most patients with mild TBI recover relatively quickly (< 3 months) [8]; however, others experience prolonged symptoms, including headaches, reduced processing speed, and attention or memory impairments [3,8].

Memory deficits are among the most common and potentially detrimental complaints among TBI patients [9–11]. Problems are associated with the hippocampus, a well-studied brain structure known especially for its contributions to memory. Earlier work has shown that the hippocampus is vulnerable to TBI and easily damaged [11–14]. Behavioral studies in rodents have proved hippocampal involvement in both working and episodic memory and that deficits occur after TBI across the severity spectrum [13]. More specifically, as measured with a standard T-maze behavior paradigm, injured mice showed impaired working memory up to 7 days post-injury, suggesting that TBI interferes with the process of memory formation [15]. Spatial memory, a subtype of episodic memory, has also been extensively studied with *in vivo* TBI models, which exhibit protracted dysfunction after mild injury [16,17].

The prevailing theory of memory describes three distinct phases – encoding, maintenance, and retrieval [18,19]. Encoding is the construction of a persistent neural representation, or memory, of an experience, maintenance entails preservation of the memory over time, and retrieval is the active process of recall or accessing the memory anew. The hippocampus is involved in all three procedures [13], but precisely how TBI perturbs these three phases remains unclear. The process of forming memories is supported by synaptic plasticity, a mechanism by which circuits are strengthened or weakened. In classical electrophysiology, such enduring, use-dependent increases in synaptic strength are encompassed by the phenomenon of long-term potentiation (LTP), or the enhancement of synaptic transmission efficiency. After TBI, several groups have demonstrated that LTP no longer occurs [20–23], especially in area CA1 of the hippocampus [21,24], suggesting plasticity impairment may underlie post-injury behavioral deficits in memory tasks. The inability to induce LTP is associated with reduced CamKII phosphorylation and synaptic protein disruption, which together represent a lower capacity for synaptic potentiation [21,25]. If LTP impairment represents a potentiation deficit and potentiation undergirds memory formation, we would anticipate encoding problems to ensue after injury. Surprisingly, there is no consensus about which phase of the memory process is most disrupted after injury.

Beyond the biological basis of memory and the disruption posed by TBI, the adaptation of microcircuit architecture through learning remains largely unaddressed in the existing literature. One tool used at the macroscale is modularity for community detection in large networks. Communities are clusters of nodes with connections to one another that facilitate performing a collaborative function [26]. The division of the brain into functional subnetworks is well-supported at the macroscale [26]. Specific to learning, one group examined how networks evolve over the course of learning through dynamic community realignment [27]. How the concepts of modularity and learning integrate in microscale circuits is unknown. While attention and learning are often studied in macroscale brain networks, there are few existing studies of learning in biologically derived microscale neural networks [28–31]. A few groups have documented how connectivity adapts with stimulation and development *in vitro* [32–34], and some models have considered learning-related input-output relationships. However, these are limited by either a lack of plasticity or specific physiological network structure. For instance, Chavlis and colleagues analyzed the effect of dendritic atrophy on pattern separation in a computational model of the dentate gyrus [28]; however, since the model does not incorporate plasticity, the results do not invoke classical potentiated learning. Examining community structure in a computational model of the hippocampus facilitates finer resolution analysis than could otherwise be obtained experimentally because we can observe the evolution of thousands of neurons over time. Furthermore, we can study the effects of an isolated mechanism of injury that has circuity-level implications. Among many possible outcomes of secondary injury sequelae, plasticity impairment can be directly linked to learning and memory dysfunction. Reports of learning-dependent network changes in this important, memory-related microcircuit are currently lacking.

In these studies, we use a model of three integrated subregions of the hippocampal formation (namely, the DG, CA3, and CA1) that comprise the classical trisynaptic circuit. The model was constructed according to known electrophysiology and anatomical connectivity data. We simulate one effect of mild TBI as STDP impairment by reducing potentiation in the circuit and establish that this deficit reduced activity and broadband power in the network. Here we extend those results by demonstrating how STDP impairment affects the structural network, focusing specifically on community organization. STDP impairment causes realignment among excitatory neurons in CA3. We also implement a learning paradigm using overlapping input patterns to study pattern separation across the hippocampal subregions. Networks with STDP impairment exhibit minor learning impairments but no pattern separation deficits, despite significant activity differences and modified community structure.

## Methods

Briefly, the model focuses on the dentate gyrus (DG), CA3, and CA1 as the primary subregions of the hippocampal formation (Fig 1A,1C). The areas follow a primarily feedforward topology with the DG sending projections to CA3 which terminates in CA1.

**Fig 1.**
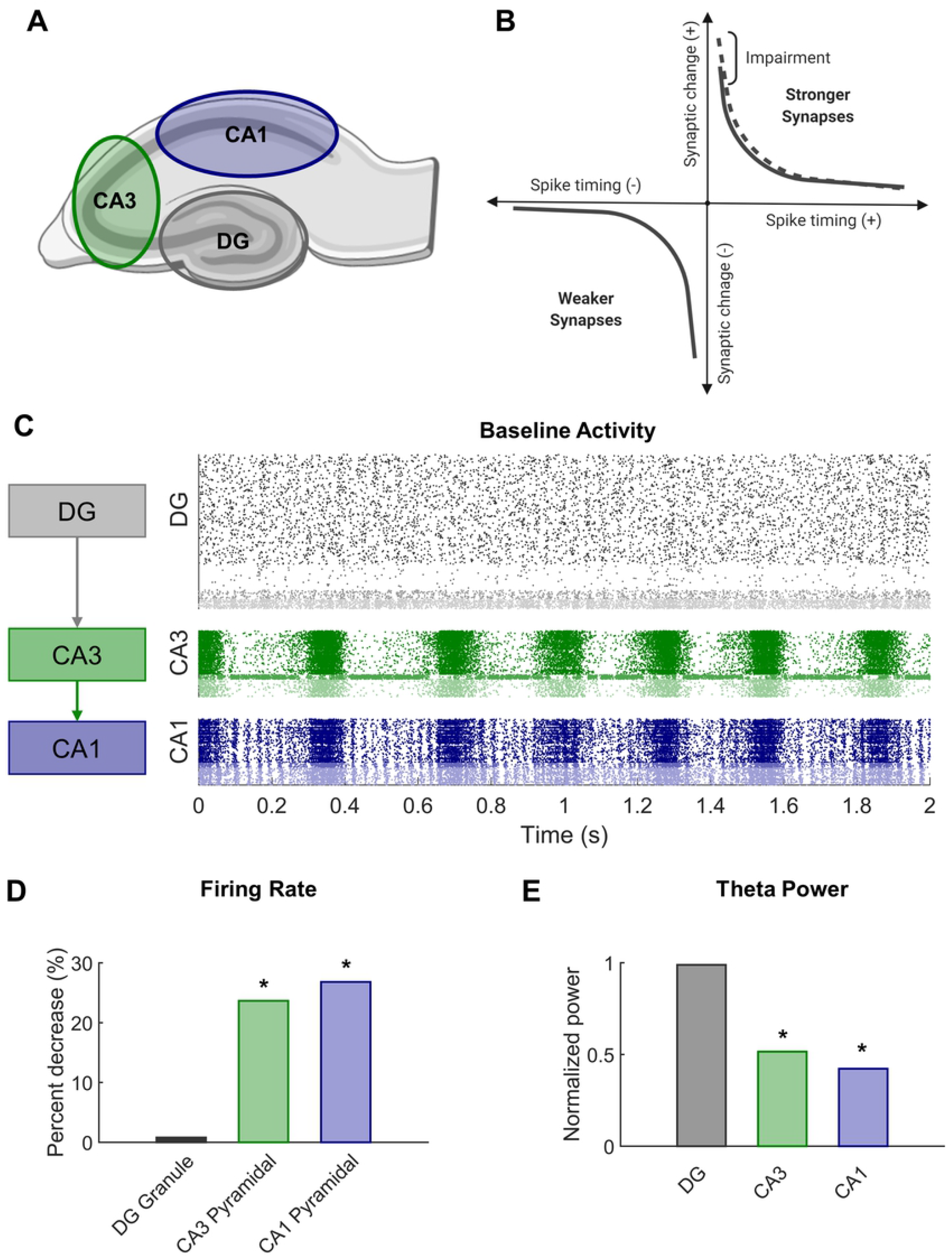
Modeling STDP impairment in a network model of the hippocampus. **(A)** The hippocampus consists of several regions connected in a predominantly feedforward topology with information passed from the DG to CA3 to CA1. These three regions are represented in the network model. **(B)** According to classical STDP, synapses between neurons with causal spikes (positive spike timing) are strengthened, but synapses between neurons with acausal spikes (negative spiking timing) are weakened. With STDP impairment, peak strengthening, or potentiation, is decreased. **(C)** At baseline, each region has a distinct pattern of firing activity. **(D)** After STDP impairment, firing rate significantly decreased in areas CA3 and CA1. **(E)** The power in the theta band, which is important for information processing and hippocampal function, also significantly decreased after injury.

### Network structure and model dynamics

The network is a system of nodes that represent neurons and edges that designate the connections between them. For each point neuron, we applied the Izhikevich integrate-and-fire neuron model, which uses the following system of differential equations to determine the spiking behavior of a neuron over time [35]:

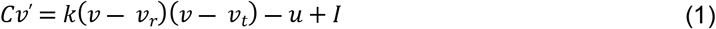

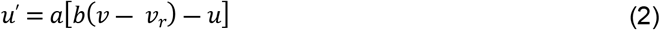

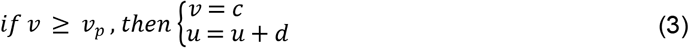

Where ***v*** is the membrane potential in millivolts (mv), and ***u*** is the recovery variable. ***C*** is the membrane capacitance (pF), ***v_r_*** is the resting membrane potential, ***v_t_*** is the threshold potential, and ***v_p_*** is the membrane potential at the peak of the spike. ***I*** is current in picoamperes (pA). The dimensionless parameters ***a**, **b**, **c***, **d**, and **k** are adjusted to represent different subtypes of neurons. The current (***I***) aggregates receptor-based ionic currents, including AMPA, NMDA, and GABA-A receptors, and 1 Hz noise input that drives the network and follows a gamma distribution (*k* = 2, θ = ½) [35–37].

There are 10 different types of neurons represented in the model across the three anatomical subregions. The dentate consists of granule cells, mossy cells, basket cells, and interneurons. Areas CA3 and CA1 each have pyramidal cells, basket cells, and interneurons with parameters specific to that subregion. Inhibitory neurons (basket cells and generic interneurons) account for approximately 10% of the neurons in each subnetwork [38–41]. The subtypes have characteristic electrophysiology and connectivity, which are represented through functional and structural features of the model, respectively. Broadly, the connectivity of the hippocampus follows a feedforward architecture. Granule cells, the principal excitatory neurons of the dentate, synapse onto CA3 neurons but have no connections to one another under physiological conditions. CA3 pyramidal cells are known to have a relatively high proportion of recurrent collaterals, but the majority of their axons project to CA1 pyramidal cells. In total, there are 8,885 neurons in the model, which converts to a scale of approximately 1:185 principal neurons in the rat hippocampus.

### Plasticity implementation and impairment

The model incorporates two primary forms of synaptic plasticity – spike-timing-dependent plasticity (STDP) and homeostatic plasticity (HSP). STDP is a form of order-dependent Hebbian learning. The process relies on precise spike timing between neurons and strengthens synapses when neurons fire causally (i.e., when the upstream neuron fires before the downstream neuron) [42]. Synaptic strengthening and weakening occur according to the following equation [43]:

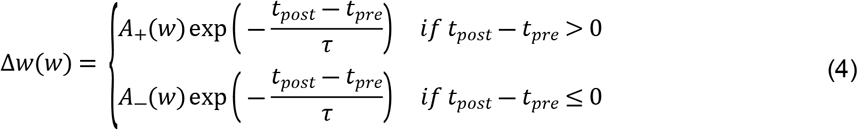

Where ***w*** is the weight of the connection between two neurons. ***A_+_*** and ***A_−_***. determine the magnitude of maximal synaptic change. The A_+_/A_-_ ratio is often biased toward strengthening and equaled 1.05 in this work [44]. ***τ*** is the plasticity time constant and was approximated as 20 ms [44]. Finally, ***t_pre_*** and ***t_post_*** are the timing of pre- and post-synaptic spikes, respectively.

Similar to previous models, plasticity applied to excitatory-to-excitatory synapses only [44]. While there are documented cases of inhibitory plasticity, inhibitory STDP is highly variable [45,46], making it difficult to implement in the model without further empirical study within this circuit. To stabilize connection weights in the network [47], we incorporated synaptic scaling, a specific form of HSP that operates at the level of individual neurons [48]. The activity of each neuron is compared to a target firing rate, and all the synapses of the neuron are modified to shift the actual firing rate closer to the target firing rate [49,50]. The following equation describes a threshold formulation of HSP adapted from [43]:

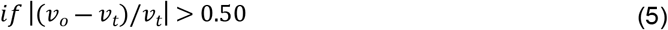

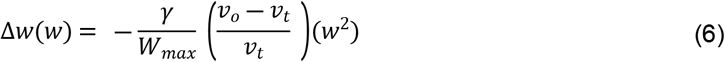

Where ***w*** is the weight of connection, ***γ*** is the dimensionless rate of change and equals 10^-8^ in these studies, ***v_o_*** is the observed firing rate, ***v_t_*** is the target firing rate, and ***W_max_*** is the maximum excitatory weight of that neuron subtype. The function has a threshold such that synaptic weights are adjusted for neurons with firing rate change greater than 50% of their target firing rate (*v_t_*) over the course of 120 s. This threshold ensures that the network continues to adapt with STDP without creating neurons with unconstrained, runaway activity.

STDP is associated with the well-studied phenomenon of long-term potentiation (LTP) observed in brain slice electrophysiology [42]. LTP describes the prolonged increase in synaptic efficacy of a circuit and is believed to support learning at the organismal level. TBI leads to deficits in spatial learning and LTP [13,20,22,23], especially within CA1 of the hippocampus [21,24]. We sought to mimic a plasticity deficit and effects of mild TBI by altering the STDP algorithm in our model. To achieve this impairment, we reduced the maximal amount of potentiation in the model by 10% (A+ = 0.9 instead of 1.0 in Equation 4 above) (Fig 1B). In our previous work, we demonstrated that this modest decrement contributed to significant decreases in firing rate and signal power in impaired networks (Fig 1D,1E). Simulations ran for 20 min without HSP to expedite synaptic settling and then 30 min with HSP. Simulations with STDP impairment were run for an additional 30 min. Analysis was performed on the final 5 min of simulation time for both baseline and impaired networks.

### Modularity analysis for community detection

Large network architectures can be partitioned into several subnetworks that perform specialized functions (Fig 2A,2B). These modules or communities generally contain densely connected nodes that are more weakly connected to other nodes outside the module. There are many methodological options for conducting community detection in networks [26]. Since our networks are directed, weighted, and signed in addition to being large (more than 3000 nodes), we required algorithms that could accommodate networks with this combination of characteristics. Modularity is one common technique used to detect the community structure of a network. Reorganizing the original matrix based on its underlying community structure takes several steps that we implemented with functions from the publicly available Brain Connectivity Toolbox [51]. Overall, we followed a procedure of modularity maximization which seeks to find the optimal network partition that maximizes the modularity quality function (Q) [26]:

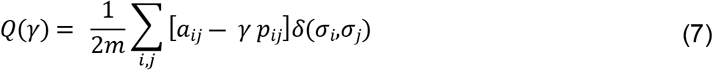

Where ***a_i,j_*** is the number of connection between modules i and j, ***p_i,j_*** is the expected number of connections between modules i and j according to a null model, *2**m*** is the total number of connections, ***γ*** is the resolution parameter, and ***δ***(*σ_i_,σ_j_*) is the Kronecker delta function.

**Fig 2.**
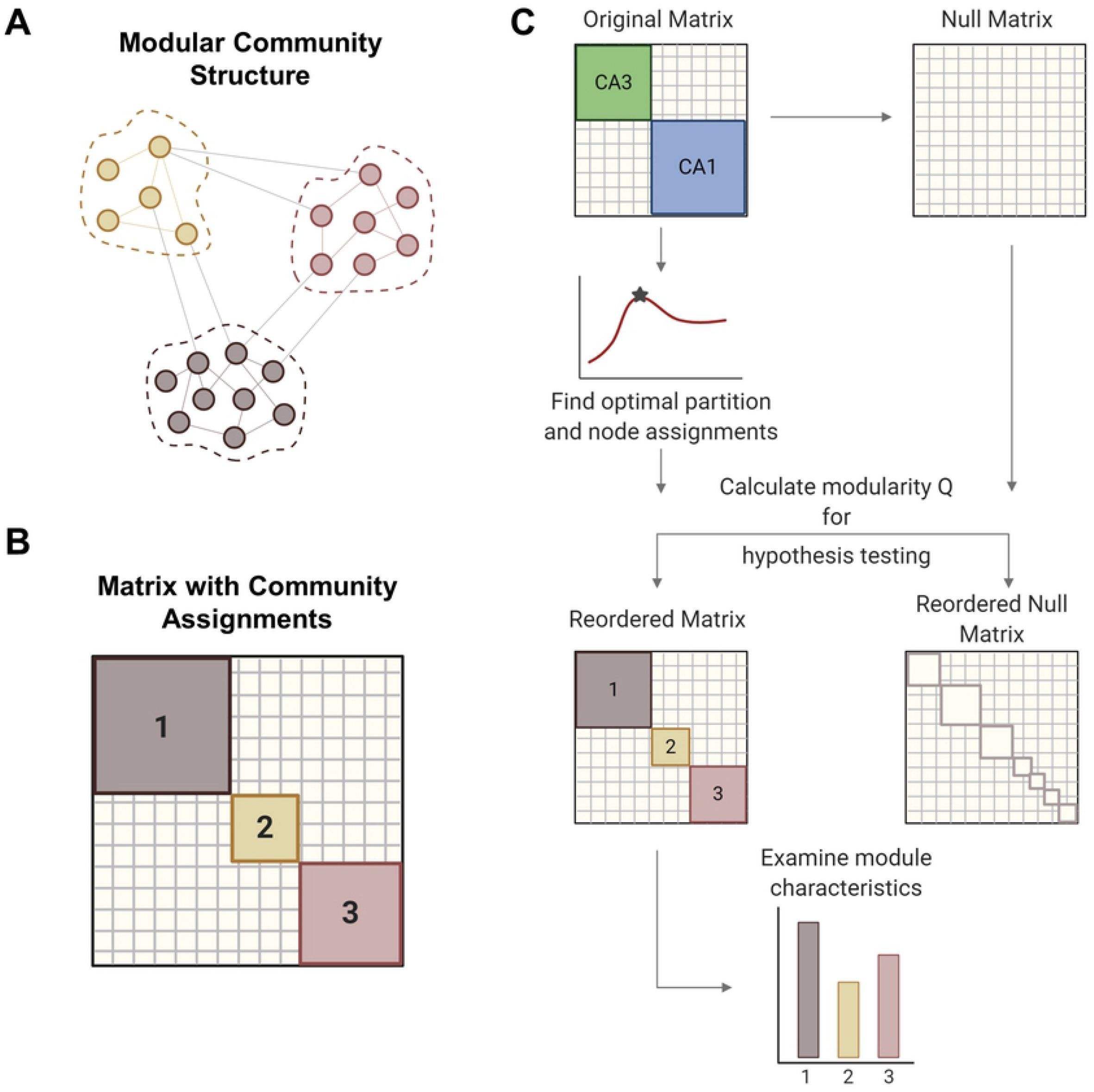
Modularity methods. **(A)** Networks can consist of interconnected modules or communities, where similar nodes are grouped with one another. **(B)** The matrix shows a network representation of community structure where neurons are grouped by module membership. **(C)** The original empirical matrix is rewired to produce the null matrix, which is a random directed graph with the same input and output degree distributions as the original matrix. The process of community detection maximizes modularity Q to find the optimal community partition. The same parameters are applied to the null matrix and module quality Q is measured for both matrices. Hypothesis testing compares the values of Q between the network of interest and the null model to verify the significance of the identified modular structure. The network is reordered based on community membership. From the reordered matrix, module size and composition can be analyzed. Created with BioRender.com.

The resolution parameter (*γ*) determines the scale of the modules that can be detected such that larger modules are detected with smaller gamma values. For hypothesis testing, a null model was generated by rewiring the original matrix while preserving the original input and output degree distributions. In gamma optimization, modularity (Q) is calculated for both experimental and null matrices across a sweep of gamma values (Fig 2C). The value of gamma that yields the largest difference in Q between the experimental and null matrices was used for subsequent steps. Gamma was optimized for minute 26 of each baseline simulation and held constant for ensuing timepoints and impaired models. With the optimized gamma parameter, we partitioned the matrix into communities many times to ensure robustness [52]. An association matrix was generated from the partition ensemble to obtain the consensus community partition. A null association matrix was also generated from a permuted partition ensemble, which is generated by permuting each column of the original partition ensemble. This null association matrix was used to threshold the experimental association matrix, thereby removing low weight connections. Consensus clustering produces an optimal partition with community assignments for each node, or neuron. Based on these assignments, the original matrix was reordered to represent the underlying community structure. We report the modularity (Q), the number and size of modules, and the composition of modules in the hippocampal networks.

### Learning and pattern separation

The hippocampus plays a key role in the broad functions of learning and memory, which depend on long-lasting, if not permanent, changes to network circuitry. These network modifications are supported by plasticity mechanisms like STDP that encode persistent responses to network stimulation. More specifically within the hippocampal formation, the dentate is known to execute the function of pattern separation, a crucial learning task in which similar incoming patterns become increasingly different from one another as they exit the network. In contrast, area CA3 with its recurrent collateral structure better supports pattern completion whereby partial pattern representations are completed as they pass through the network.

Although there are many ways to test learning in a neural network, given the size of our networks (> 8000 nodes), an unsupervised learning algorithm was preferable to a supervised approach, so we evaluated learning with a similar method to our previous work [31]. To summarize this method, we applied two protocols to assess learning. During *training*, the network was stimulated and able to adapt with plasticity to encode responses to periodic input over 30 minutes. During *testing*, static networks were stimulated for 6 minutes. Networks were tested before and after training to determine how training modified the network response.

The networks were first settled as described previously for 30 min of simulation time with 1 Hz noise and then trained with exogenous 1 Hz stimulus. For each of two patterns, we simultaneously stimulated a set of 200 input neurons in the DG and measured the response in all three subregions. The input patterns overlapped by 50% with 100 neurons that were common to both patterns and 100 neurons that were unique to each stimulus. The simulation ran for an additional 30 min with 1 Hz noise and 1 Hz stimulation of each pattern to encode the activity response before the network was tested. The response was measured in the 200-ms epoch immediately following stimulation of either pattern 1 or pattern 2.

Since learning is defined by training-dependent changes in network activity, we tested the response of *untrained* and *trained* networks to the two input patterns in order to determine which neurons augmented their activity after the training period. The activity of each neuron post-training was normalized by its activity before training to account for neurons with inherently high activity. The 200 neurons that increased their firing the most from untrained levels comprised the desired, *target* response. The remaining neurons made up the *off-target* response where increases in activity are undesirable. Thus, the response for each subregion consists of a target component of 200 neurons that respond maximally to the stimulus and an off-target component of the remaining principal neurons. The signal-to-noise ratio was measured as the ratio of the target to off-target response. Finally, this paradigm was repeated for two training conditions. One set of networks was trained under baseline conditions, and another set of networks underwent training with STDP impairment to test whether reduced potentiation interferes with the ability to encode patterned responses.

To evaluate pattern separation across the subregions of the network, we turned to several additional metrics. First, we examined the extent to which the target output populations from patterns 1 and 2 differed by calculating the percent overlap among the two populations for each network. More formally, we measured the change in population distance via the Hamming distance, which calculates the proportion of positions that differ between two binary vectors:

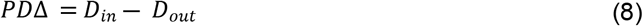

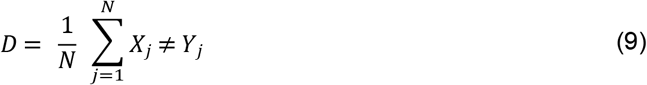

Where ***PDΔ*** is the change in population distance. ***D_in_*** and ***D_out_*** are the Hamming distance between the input and output patterns, respectively. ***X_j_*** and ***Y_j_*** are binary vectors representing patterns 1 and 2, and ***N*** is the length of the binary vectors. By this metric, identical vectors have a Hamming distance of 0 while two unique vectors have a Hamming distance of 1. If a network performs pattern separation, the Hamming distance of two input populations will be greater than that of the corresponding output populations [28]. If PDΔ > 0, the network performs pattern separation. If PDΔ < 0, the output patterns are more similar than the input patterns. A second feature of pattern separation accounts for rate differences between the output patterns [28]. For this analysis, we focused on the target neurons that were common responders to both patterns and measured the mean Spearman distance between the pattern 1 and pattern 2 responses of common neurons. The Spearman distance (SD) is calculated as one minus the Spearman rank correlation between two vectors:

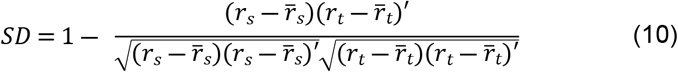

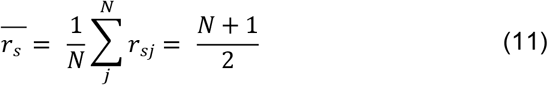

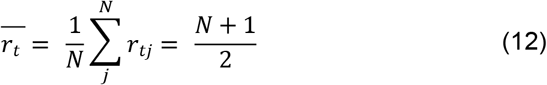

Where ***r_s_*** and ***r_t_*** are the rank vectors of ***x_s_*** and ***x_t_***, representing the normalized rate response pattern 1 and pattern 2, respectively. ***N*** is the length of the vectors and number of common neurons between patterns 1 and 2.

### Statistical analysis

For statistical comparisons between baseline networks and rewired, null models, we used Student’s t-test. To compare baseline and impaired networks, we applied a paired Student’s t-test with Bonferroni correction to determine significance for cases of multiple comparison. Statistical testing also included repeated measures ANOVA with Tukey-Kramer post-hoc test for comparisons with multiple timepoints.

## Results

### Modularity in baseline networks

For modularity analysis, we narrowed our focus to areas CA3 and CA1 due to network size and because these two subregions displayed the largest injury effects in our preliminary analysis of functional changes after impairment (Fig 3A). To establish whether the hippocampal networks had detectable community structure (Fig 3B), we compared them to null models generated by rewiring the connections of the original matrix while preserving the input and output degree distributions. We found that the number of modules was significantly lower in the hippocampal model matrices than in the randomized networks, indicating that empirical communities are more integrated than predicted by random models (Baseline hippocampal: 6 ± 0.5 vs. Randomized: 24.9 ± 1.5; Student’s t-test; p < 1e-10) (Fig 3C). As expected, modularity (Q) was significantly higher in experimental baseline networks than in randomized controls (Baseline hippocampal: 0.269 ± 0.002 vs. Randomized: 0.089 ± 0.001; Student’s t-test; p < 1e-10) (Fig 3D). High values of Q mean that the detected communities have higher internal connectivity than predicted by chance. Together, these results confirm that the hippocampal networks have significant modular structure as compared to null models. Furthermore, we evaluated modularity Q for the last 5 min of simulation time at baseline and found no change in Q over time (One-way ANOVA; F-statistic = 0.08; p > 0.5) (Fig 3E). Therefore, we used the final connectivity matrices (from min 30) to compare baseline and impaired networks in subsequent analysis.

**Fig 3.**
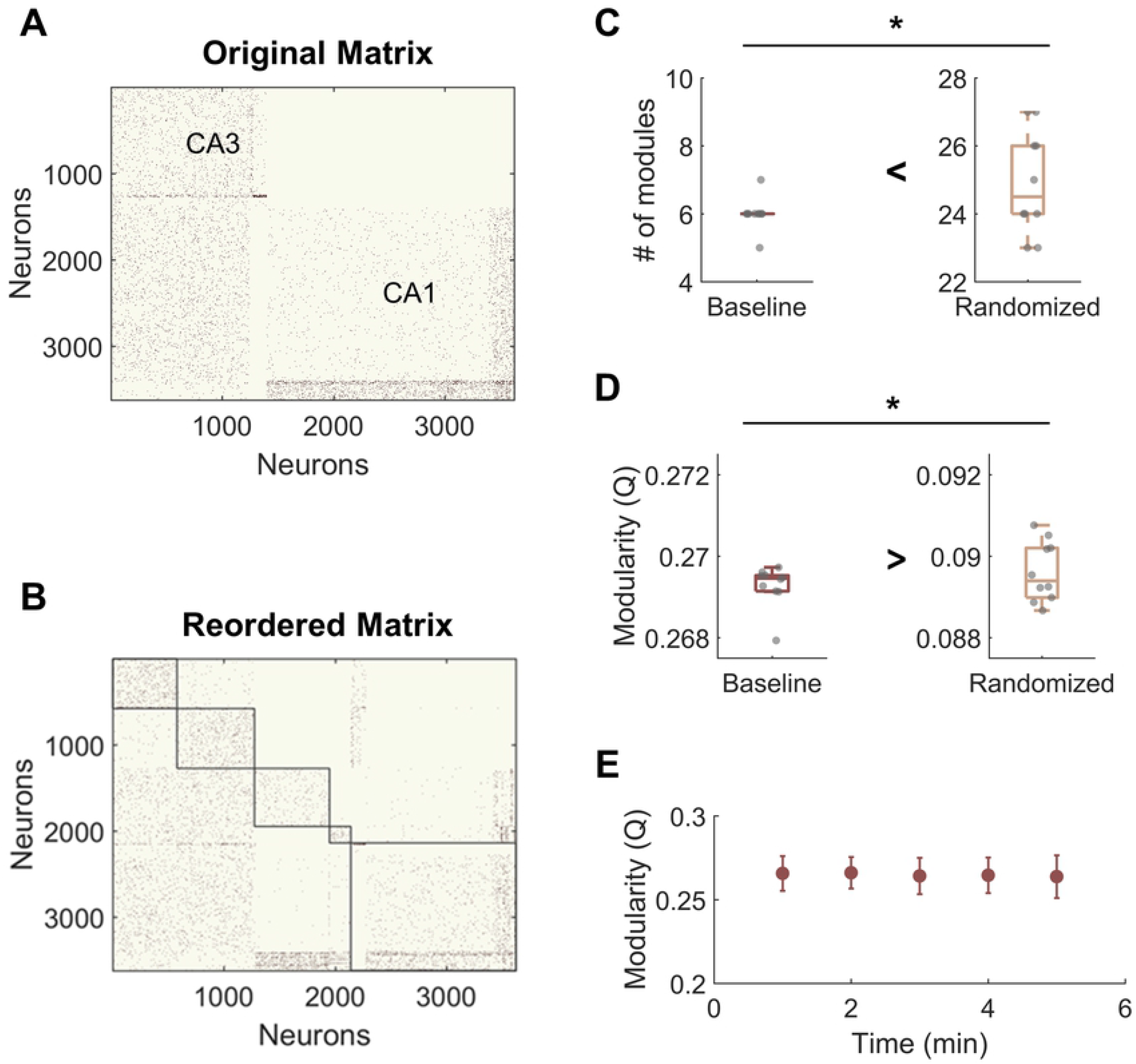
Hippocampal model networks have significant community structure compared randomized control networks. **(A)** A representative baseline network organized by anatomical structure (CA3 vs. CA1). **(B)** A representative network reorganized by module. **(C)** The number of modules is significantly higher in the randomized networks than at baseline (p < 1e-5). **(D)** Modularity, Q, is significantly lower for randomized networks (p < 1e-5). Randomized controls rewired connections in the original network while preserving the degree distribution. **(E)** There was no significant change in modularity over time at baseline.

### Effects of STDP impairment on community structure

We next compared the community structure of baseline networks with that of STDP impaired networks (Fig 4A,4B). Models with STDP impairment ran for an additional 30 minutes, and the ending connectivity was compared to the pre-injury connectivity using the same modularity algorithm and holding gamma constant. Modularity Q decreased significantly after plasticity impairment (Baseline: 0.26 ± 0.01 vs. STDP Impaired: 0.24 ± 0.02; paired Student’s t-test; p < 0.01) (Fig 4D). However, the number of modules did not differ (Baseline: 5.0 ± 1.0 vs. STDP Impaired: 5.3 ± 1.3; Student’s t-test; p > 0.1) (Fig 4E). While the average number of modules per network remained the same, we did identify trends in the sizes of modules after injury. Modules derived from networks with STDP impairment were more likely to fall at the extreme ends of the size range (Fig 4C). In particular, there are more small communities below a size of 250 nodes. On a network level, the size range between the largest and smallest module of each network increased after STDP impairment, reflecting the evolution of these smaller communities (Baseline: 1129 ± 333 vs. STDP Impaired: 1439 ± 332; Student’s t-test; p < 0.05) (Fig 4F).

**Fig 4.**
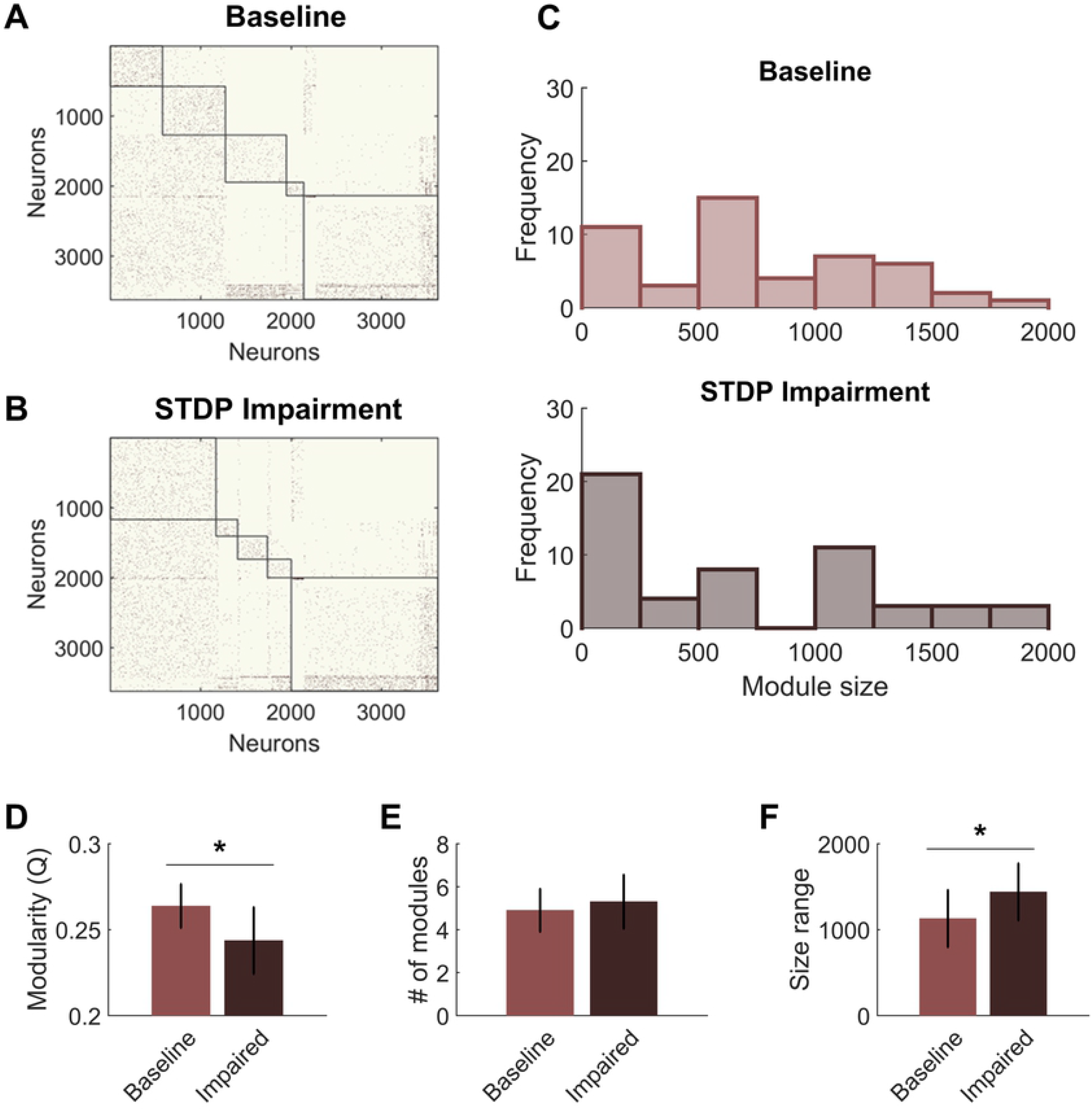
STDP impairment decreases modularity in the CA3-CA1 network. **(A)** A representative network organized by community assignment shows 5 modules at baseline. **(B)** The same representative network has 5 communities after STDP impairment, but individual node assignments can change resulting in different module size characteristics. **(C)** Histograms of module size across all 10 networks show that there are more modules at the extreme ends of the size range after STDP impairment. **(D)** Module quality Q decreased significantly with injury (p < 0.01). **(E)** The average number of modules per network did not change after injury. **(F)** The range of module size increased significantly after injury (p < 0.05).

The shifts in module size suggested a broader realignment of neurons among existing communities, and we further hypothesized that the detected community structure might reflect the anatomical designations of the hippocampal circuitry. Therefore, we analyzed the neuron subtype composition of each module for both baseline and impaired networks. Each module was characterized based on the percentage of neurons from CA3 vs. CA1 and the percentage of inhibitory neurons. We found that excitatory neurons from CA3 tended to segregate into their own communities (Fig 5A). The remaining communities contained most of the CA1 excitatory neurons (pyramidal cells) as well as inhibitory neurons from both CA3 and CA1. Accordingly, we identified a significant relationship between the percentage of inhibitory neurons in the module and the percentage of CA1 neurons. As the inhibitory percentage increased, the percentage of CA1 neurons decreased, indicating that these additional inhibitory neurons were anatomically derived from CA3 (Y = 0.40X + 0.005; linear regression; R^2^ = 0.75; p < 1e-5). After STDP impairment, we found that CA3 excitatory neurons continued to form separate communities; however, the relationship between the percentage of inhibitory neurons and CA1 neurons disappeared (Fig 5B). This occurs due to the appearance of many small modules that contain excitatory neurons from both CA1 and CA3. Most likely, some neurons from the CA3 excitatory modules realign with excitatory neurons from CA1 to form these small communities.

**Fig 5.**
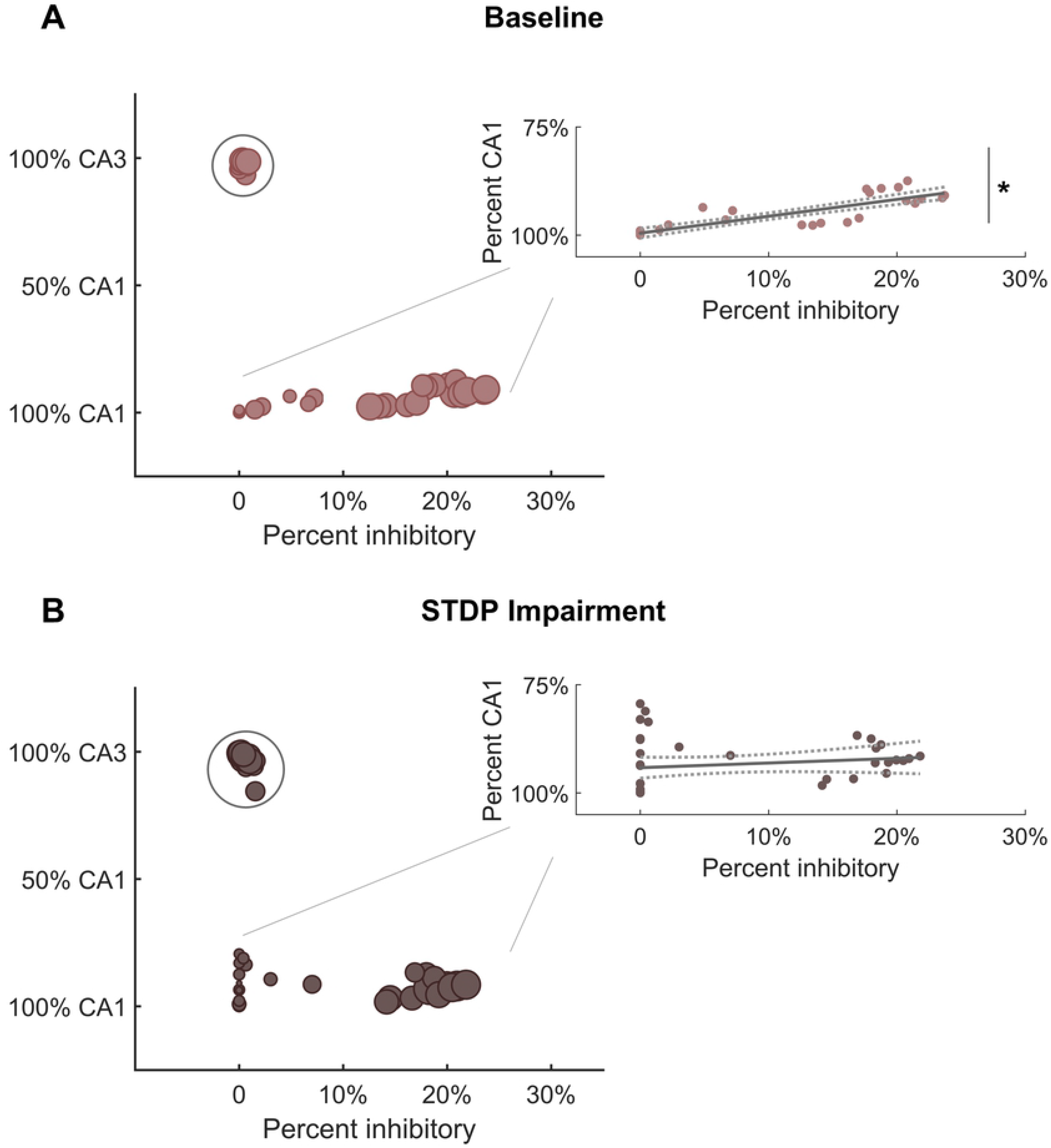
Module characterization by underlying neuron type reflect hippocampal anatomy. **(A)** At baseline, one subgroup of modules is comprised primarily of CA3 excitatory neurons (within circle). Predominantly CA1 modules contain most of the inhibitory neurons from both CA3 and CA1. Therefore, there is a significant relationship between the percentage of inhibitory neurons and the percentage of CA1 neurons in these modules (inset) (R2 = 0.75; linear hypothesis test; p < 1e-5). As the percentage of inhibitory neurons increases, the percentage of CA1 neurons decreases (inset). **(B)** After STDP impairment, there remains a subgroup of modules comprised of CA3 excitatory neurons (within circle). However, a new subgroup of small modules develops. These are made up of excitatory neurons from both CA1 and CA3. The appearance of these small excitatory modules eliminates the relationship between inhibitory tone and the percentage of CA1 neurons (inset) (R2 = 0.03; linear hypothesis test; p > 0.1).

### Pattern separation in baseline and impaired networks

Learning and memory are crucial hippocampal functions supported by synaptic potentiation. As a mechanism of synaptic weight modification, STDP facilitates use-dependent circuit adaptation. To test whether and how STDP impairment affects higher-level network functions, we implemented a method of unsupervised learning characterized by training-dependent changes in neural activity (Fig 6A). Baseline networks were trained with STDP impairment or under control conditions. During *training*, two overlapping sets of 200 neurons in the DG were stimulated in addition to receiving baseline noise input. The two stimulus patterns were interleaved and stimulated at 1 Hz. During *testing*, the same two input patterns were activated in a static network. Networks were tested before and after training to determine the relative change in firing rate on a neuron basis. Not including those neurons stimulated with input patterns, the rest of the principal excitatory neurons in the network were divided into two groups of responders. For each subregion (DG, CA3, CA1), those that increased their spiking activity the most were termed target neurons, and the remainder were classified as off-target neurons. Target and off-target neurons were not identified *a priori* but rather based on their response to the training paradigm.

**Fig 6.**
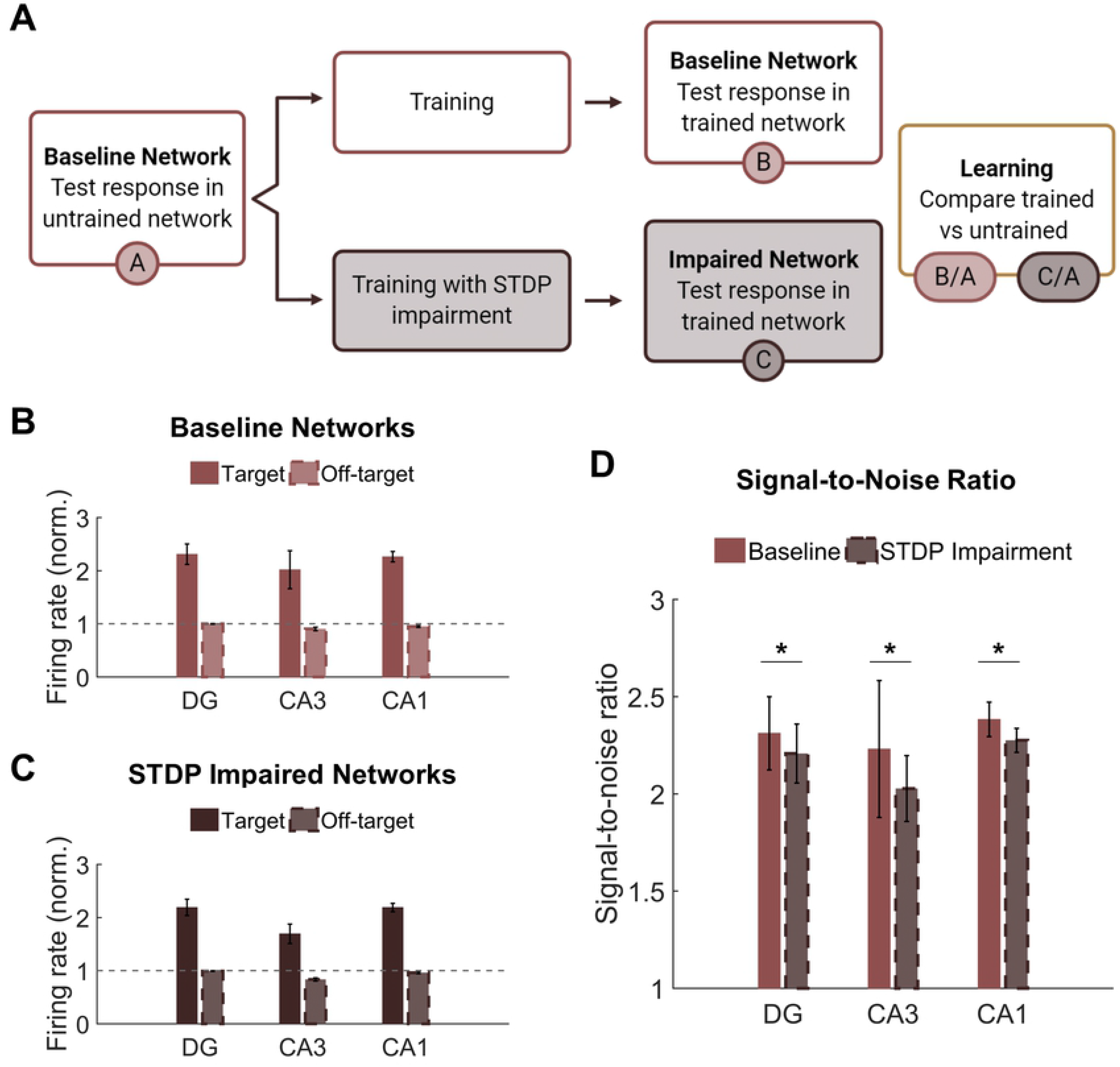
Networks successfully encode patterned responses although STDP impairment decreases the signal-to-noise ratio. **(A)** Training consisted of stimulating sets of 200 neurons in the DG. Baseline networks were trained once with STDP impairment and once under control STDP conditions. Networks were tested before and after training to compare the activity response in each region. **(B)** Firing rates after training are normalized by the response to stimulation in the untrained network. The gray dashed line is the reference point for activity in untrained baseline networks. The activity of target neurons increases significantly from baseline while the average activity of off-target neurons remains the same or decreases. **(C)** Networks with STDP impairment exhibit the same paradigm as baseline networks with higher activity in target neurons than in off-target neurons. **(D)** The signal-to-noise ratio (on-target divided by off-target response) decreases significantly after injury in each region (paired Student’s t-test, p < 0.02 with significance determined by Bonferroni correction).

Although we hypothesized that limiting potentiation would interfere with the encoding phase of memory, we found that both baseline and STDP impaired networks were capable of encoding conditioned responses to input stimulation. The target neurons had significantly higher average normalized firing rate than their off-target counterparts across all three subregions and both conditions (Student’s t-test; p < 1e-5 for all conditions) (Fig 6B,6C). We also computed the signal-to-noise ratio (SNR) as the target activity divided by the off-target activity and found that STDP impaired networks expressed lower SNR in all three subregions of the hippocampus with the most significant change in CA1 (Paired Student’s t-test; p < 0.02 for all subregions) (Fig 6D). This decrease in SNR appears primarily driven by a decrease in firing among target neurons. Although significant, the magnitude of the difference was modest.

Thus far in our analysis of the learning paradigm, we focused only on the magnitude of the output; however, we also investigated whether the response to each pattern differed. Given the observed decrease in SNR among the responder neurons, we sought to determine whether this decrease affected the ability of the circuit to perform pattern separation by discriminating between the two overlapping input patterns. Successful pattern separation requires that the output patterns be more different than the input patterns (Fig 7A). Accordingly, we evaluated the amount of overlap between the groups of target neurons for each pattern, finding that the mean percentage of overlap was 16% and 13% for the DG and CA1, respectively (Fig 7B). This is well below the 50% overlap of the input patterns, indicating strong pattern separation. Interestingly, the percentage of overlap among target neurons from CA3 was 48% on average (Fig 7B), so this area did not execute pattern separation. This is most likely attributable to the recurrent collaterals in CA3 that putatively make the area uniquely adept at pattern completion, the ability to complete an output response based on partial input information. Due to the limited ability to potentiate synapses, we hypothesized that STDP impairment would limit the ability to encode unique output patterns. However, we found that the percentage of overlap did not decrease in networks that were trained with impairment. In fact, the change in population distance between the input and output populations increased in the DG and CA3 of impaired networks, suggesting that pattern separation was more successful in these subregions (DG: 0.18 ± 0.06 vs. 0.24 ± 0.04; paired Student’s t-test; p < 0.001. CA3: −0.15 ± 0.04 vs. −0.03 ± 0.05; p < 0.001. CA1: 0.21 ± 0.03 vs. 0.21 ± 0.02; p > 0.1 for CA1) (Fig 7C).

**Fig 7.**
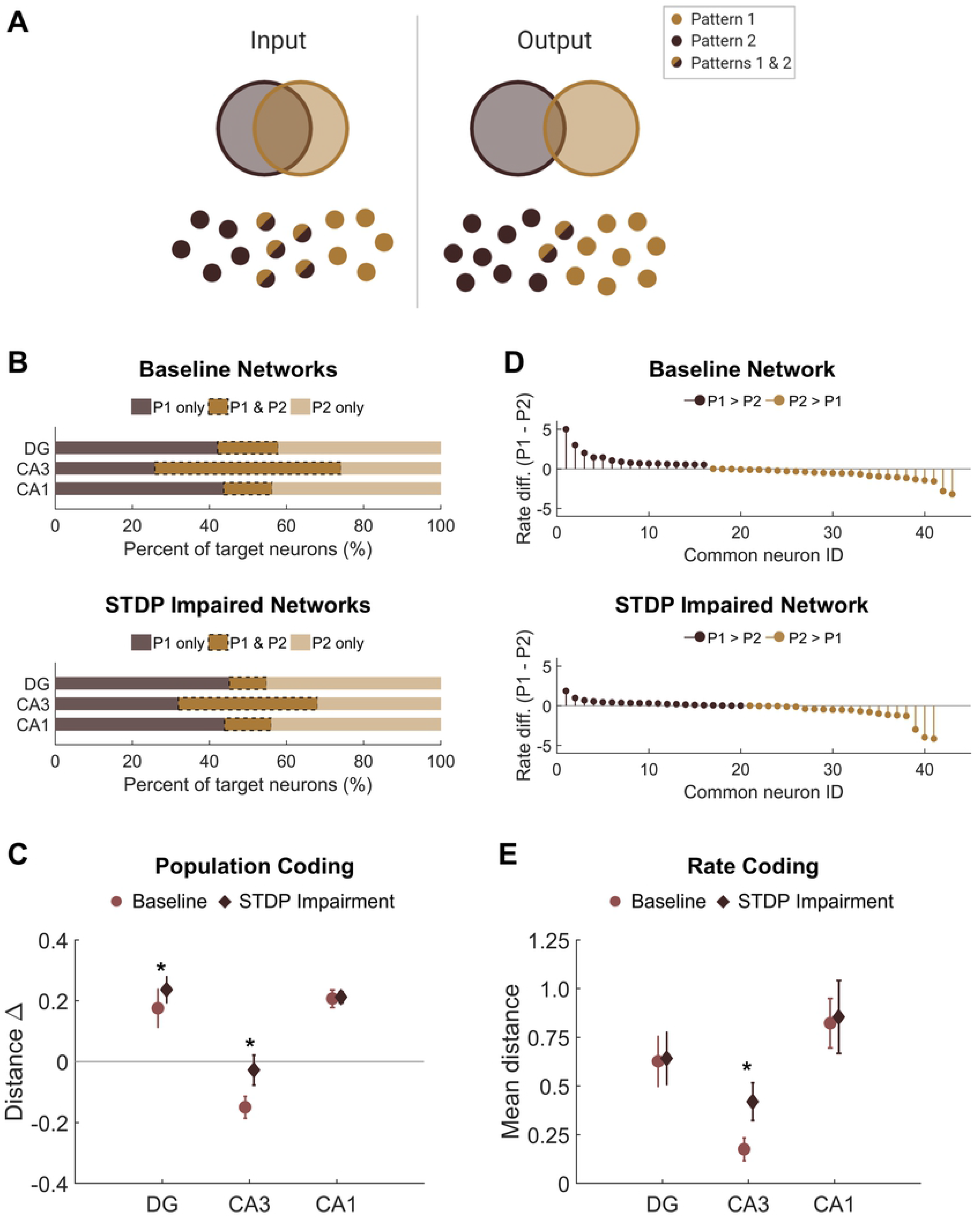
There is no pattern separation deficit in circuits with STDP impairment. **(A)** Pattern separation occurs when the output patterns differ more than the input patterns do. In this study, we stimulated two patterns with 50% overlap in the population of input neurons. **(B)** For each region, the output populations consisted of 200 target neurons for each pattern. The percent overlap in baseline networks was below 20% for the DG and CA1. Similar to baseline networks, STDP impaired networks had low percentage overlap in the DG and CA1 with higher overlap in area CA3. **(C)** The difference between the Hamming distance of the input population and the output population measures pattern separation where a higher value indicates greater pattern separation. With STDP impairment, the distance between output populations was greater in the DG and CA3 than at baseline (paired Student’s t-test, p < 0.02 with significance determined by Bonferroni correction). **(D)** The rate difference between common neurons shows that common neurons responded preferentially to one pattern or the other. Common target neurons from the DG in one representative network are shown. P1 = pattern 1; P2 = pattern 2. **(E)** The distance between the rate response to pattern 1 vs. pattern 2 was computed as the Spearman distance. The rate distance for CA3 outputs was significantly different between baseline and STDP impaired networks (paired Student’s t-test, p < 1e-5).

In addition to distinct populations of responsive neurons, rate coding is another attribute of pattern separation [28]. Since most of the neurons in the target populations were unique to one pattern or the other, we were interested in the neurons that activated with both patterns and whether these common neurons responded preferentially to either pattern. We calculated the normalized rate difference between pattern 1 and pattern 2 activity for all common neurons (Fig 7D). To compute the distance between the response vectors, we evaluated the mean Spearman distance across networks. We found that the only subregion to show a significant change after STDP impairment was CA3, but there were no significant differences in rate coding among common neurons of the DG or CA1 (Paired Student’s t-test with Bonferroni correction for multiple comparisons; p < 1e-5 for CA3) (Fig 7E). Therefore, although STDP impairment reduced the total SNR, rate coding was still effective for pattern separation among common responder neurons. While no deficits were observed in population- or rate-based analyses of pattern separation in these circuits, these results do not preclude the possibility that there may be subtle differences in temporal coding based on specific spike timing.

### Nodal flexibility in target neurons

Finally, we assessed modularity in trained baseline and STDP impaired networks. Similar to untrained impaired networks, trained circuits with STDP impairment had lower modularity than untrained baseline networks (Repeated measures ANOVA with Tukey-Kramer post-hoc for multiple comparisons; p < 0.05) (Fig 8A). Trained baseline networks did not significantly differ from either untrained baseline or STDP impaired networks (Fig 8A). After verifying community structure in trained networks, we investigated how community affiliations changed over time. To do so, we applied the concepts of ‘flexibility’ and ‘promiscuity’ (Fig 8B). As it relates to network theory, flexibility describes whether nodes change their community affiliation at different time points. Nodes with high flexibility frequently associate with different modules. Promiscuity is a related yet distinct concept that quantitatively captures whether nodes associate with several unique modules or only a few. A highly flexible node could have low promiscuity if it shifts between only two unique communities. We analyzed flexibility and promiscuity in the aggregate target and off-target populations of CA3 and CA1. Target neurons, which increased their activity the most after training, were most likely to fall in the highest or lowest quintiles of the flexibility distribution (Fig 8C). These neurons also had low promiscuity, indicating that changes in community assignment included few unique modules (Fig 8D). Together, these results suggest that target output neurons have comparatively stable community affiliations since even those that were flexible were associated with lower promiscuity. In contrast, off-target neurons fell relatively evenly into the flexibility and promiscuity quintiles (Fig 8C,8D). We found no significant differences in these properties after STDP impairment.

**Fig 8.**
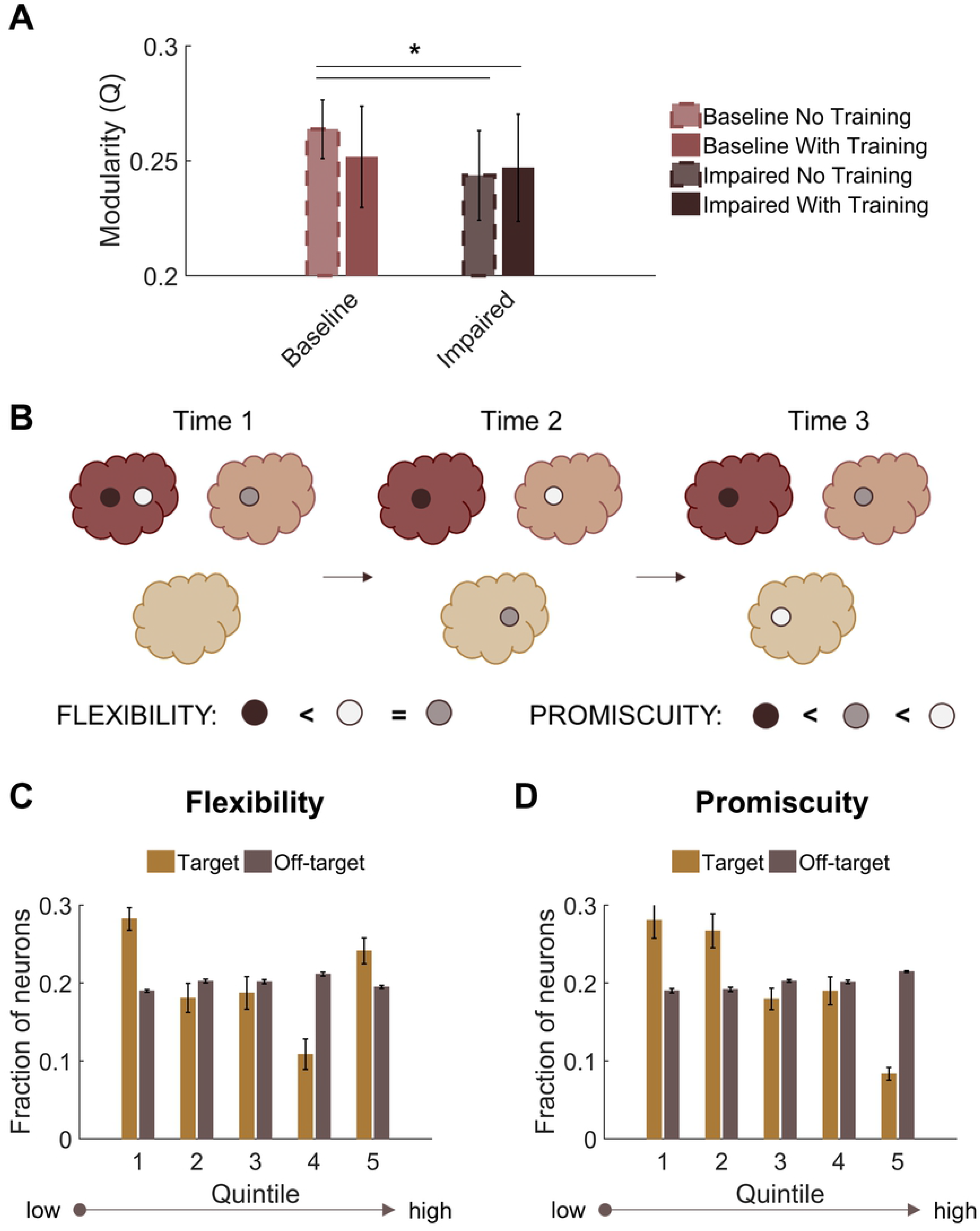
Target output neurons have low promiscuity among network communities. **(A)** Both trained and untrained networks with STDP impairment have lower modularity than untrained baseline networks. **(B)** Neurons that change their community affiliation frequently have high flexibility. If their affiliation shifts between unique communities, those neurons also have high promiscuity. **(C)** Target neurons are more likely to fall in the first or fifth quintiles of the flexibility distribution. **(D)** Target neurons have low promiscuity, most likely falling into the first two quintiles of the distribution.

## Discussion

In these studies, we examined the community structure of a neuronal network model of the hippocampus. At baseline, we found that the CA3-CA1 networks displayed significant modular structure in which excitatory neurons from CA3 (pyramidal cells) reliably segregated into distinct communities. The remaining neurons, including CA1 pyramidal cells and inhibitory neurons from both subregions, formed separate modules. After STDP impairment, modularity decreased significantly, and more small modules appeared. With their small, spurious nature, these modules are purportedly less functionally well-defined. We then trained the networks with an unsupervised learning algorithm to test the critical function of pattern separation across the subregions of the circuit. In the learning process, we identified a critical group of target neurons that showed the largest rate-dependent training effect. STDP impairment during the encoding phase of pattern acquisition reduced the magnitude of the learning effect; however, impaired networks executed pattern separation successfully as analyzed with both population- and rate-based coding. Finally, we found that target neurons had a unique modularity-derived profile characterized by low nodal promiscuity, which indicates that these target neurons were relatively stable and affiliated with few unique network communities. In comparison, off-target neurons followed more homogenous flexibility and promiscuity distributions.

There are several limitations to the current studies that influence the interpretation of this work. Fundamentally, the hippocampal model is not full-scale and contains a limited number of cell and receptor types. It does not have lamellar structure or complex geometry; however, the synaptic connectivity is faithful to the literature and the most important attribute for the network-based analysis presented here. We use a point neuron model of Izhikevich integrate-and-fire neurons that is more phenomenological than other, more biophysical neuron models. This drawback is balanced by high computational efficiency, which enabled the development of a large network model of the hippocampus, and by extensive use and validation of cell-specific spike timing across different neuron types [35,53–56]. Our simulation of STDP impairment as a consequence of mild TBI is also a limitation of these studies. Despite the prevalence of learning and memory deficits after TBI, there is not extensive literature surrounding plasticity impairment. Beyond inhibiting long-term potentiation in hippocampal circuitry, injury reduces CaMKII expression and synaptic protein assemblies, thereby impeding synaptic strengthening. In contrast, long-term depression generally remains intact in damaged hippocampal slices. Accordingly, we modeled these effects as a biased decrement in potentiation only. The change was modest, consisting of a mere 10% decrease in maximal strengthening, to ensure that network activity did not collapse at baseline. However, given our results that STDP impairment did not have a strong negative effect on learning and pattern separation, additional injury mechanisms should be explored in future work. Since damage is known to cause pattern separation deficits in both animals and humans [13,57,58], our results suggest that some additional mechanism beyond STDP impairment must contribute to those effects. Upcoming modeling studies might also examine the interplay between different plasticity algorithms since a stronger homeostatic mechanism might compensate for larger decreases in STDP-related potentiation, thereby preserving baseline untrained activity levels while exposing larger learning deficits.

Lastly, we implemented an unsupervised learning paradigm, which makes no *a priori* designation between desired and undesired responses. For each of two patterns, we stimulated a subset of 200 neurons in the dentate and identified the most responsive neurons in all three hippocampal subregions based on their normalized firing rate. We also implemented training and STDP impairment at the same time to hold the runtime constant between impaired and control networks. Yet, we could also consider training networks that had already adapted to STDP impairment controlled. It is possible that training mitigated the effects of injury and that networks with ingrained diminished activity are less responsive to training. Although this unsupervised method of network learning cannot address complex temporal coding, it has several advantages. Since it is a computationally efficient post-hoc algorithm without prior topological assumptions, it could be applied with spiking data of this size and density. It also exploits our incorporation of use-dependent plasticity (STDP) as one of the major advances in a model of this size and biological fidelity. Therefore, this method constituted a reasonable biological proxy despite its unsupervised nature. One popular alternative in biologically inspired neural networks is the detection of polychronous neural groups, which is better adapted to handling many neural groups and memory traces and evaluating the maximal amount of information that might be stored in a given circuit. While the original algorithm requires computationally expensive, brute-force computations, some groups are developing more efficient alternatives inspired by the field of machine learning [59,60]. These approaches might offer an opportunity to extend these results with a quantification of the information storage capacity of this hippocampal circuit.

Modularity is a useful framework for assessing the architectural organization of a network. Large networks often consist of several smaller subnetworks that are more densely connected internally than they are externally [26]. This partitioned organization is posited to support faster, more efficient processing by facilitating functional compartmentalization [26,61]. By reducing the energy requirements for network-wide modifications, a more modular structure is also a more adaptable one [27,62]. In the present study, we identify post-injury modularity reduction, which may constitute an adaptation that increases integration between communities to support overall activity levels. This result further suggests that potentiation supports baseline segregation in the hippocampal circuit. Although there is also evidence of the opposite [63,64], previous findings that TBI reduces modularity in functional brain networks correlate with persistent post-concussive syndrome [62,65]. In addition, more modular structure was predictive of better training outcomes after injury [62]; this may occur because lower energy costs are associated with adaptation in more highly segregated networks. The variable response (increase vs. decrease) may relate to individual heterogeneity, other measures of network-wide integration, or whether the network is still in a state of active adaptation. Aside from analyzing the microcircuit scale, differences between our results and others may be attributable to our focus on structural, instead of functional, connectivity. Since functional connectivity is dynamic and malleable while structural connectivity is more stable as a reflection of the underlying neural anatomy, it is possible that the training effect is larger for functional networks. Ultimately, the demonstrated effects in structurally well-defined microcircuits corroborate the idea that modularity may be a useful (bio)marker of intervention-dependent network plasticity [66].

With a modest amount decrement in STDP-related potentiation, networks could still learn and execute pattern separation. In another recent study from our group, we tested the circuit-level consequences of NMDA receptor damage, which increases network activity, in a generic circuit with a similar learning paradigm [31]. Since injury-induced, elevated activity obscured the training effect, we found the most detrimental outcome of NMDA receptor dysfunction occurred during recall of previously trained patterns but also tested injury during different phases of memory [31]. Here, we exclusively tested STDP impairment during the encoding stage. In our preliminary work analyzing impaired networks without exogenous stimulation, we found significant declines in firing rate and signal power. Based on those effects, we expect the maintenance phase would also challenge STDP impaired networks, which would likely lose the entrained response more quickly as overall activity decreases without exogenous stimulation. This idea is supported by behavior studies that find injured animals perform the task successfully if tested quickly after training but not if the time between testing and training is longer [16]. If we integrated both STDP impairment and NMDA receptor damage simultaneously, we expect that STDP impairment might enhance pattern recall because the two mechanisms have opposing influence on network activity. Alternatively, as trauma-induced changes to NMDAR physiology will disappear when receptors are replaced hours after injury [67–69] and plasticity impairments may persist for days after injury, one might expect an acute early impairment in the retrograde memory [70–72] with a longer lasting impairment in memory acquisition [16,73,74].

As a measure of encoding, pattern separation conveys the capacity to distinguish similar events and contexts; therefore, this function underpins general learning abilities. TBI causes behavioral deficits in spatial memory and spatial object recognition in animal models of injury [13,16,75]. These trained behaviors depend on discrimination between similar experiences. Recent findings also demonstrate that injury impairs pattern separation in humans [57,58]. The dentate is traditionally the primary focus of studies on hippocampal pattern separation because its intrinsic properties of high inhibition and parallel circuity intuitively support this filter function; however, there is growing evidence that other subregions (CA3 and CA1) also facilitate pattern separation. In fact, temporary CA1 lesions impair pattern separation in humans [58]. Since CA1 relays information processed by the hippocampus to neocortical brain regions [76], the area clearly plays an important role in the wider circuitry, making it an intuitively important subregion. For these reasons, it is interesting that in our work changes in the DG and CA3 appear to compensate for one another because the population- and rate-based output distances measured from CA1 do not differ. These results suggest that the output patterns transduced by CA1 are essentially the same and that the network adapts to maintain that final output. While one might predict a larger effect of STDP impairment on pattern separation, these subtle differences are an intuitive extension of our collective results. With our preliminary functional analysis in networks without learning, we found that the DG was remarkably robust after STDP impairment. Specifically, the average firing rate and signal power did not decrease significantly. Given its intrinsically low rates of activity, the DG is more resilient to minor changes in STDP. Others have found that deficits in pattern separation are associated with hyperexcitability and elevated activity in the DG [28,77,78], which increases activation and thereby reduces the capacity of the filter function. In general, STDP impairment reduces synaptic weights in the network, making it more difficult to activate. Although its impact on learning is more indirect, NMDA receptor damage or inhibitory cell degeneration might have outsized influence on pattern separation because these mechanisms would increase spurious noise in the output patterns. Finally, our analysis in this work focused on population and rate coding; however, we cannot exclude the importance of temporal coding because it is possible that the spike timing changes while the activity rate remains stable. Indeed, previous results from our group indicate that networks adapt to preserve firing rate first as other measures of spike timing exhibit longer lasting changes after neurodegeneration [37].

One natural extension of this work is prospective training or other interventions designed to facilitate active recovery in damaged networks. For instance, a stimulation protocol that could restore activity in a damaged network would be of interest for rehabilitation [13,79], and certain types of stimulation (frequencies, magnitudes, etc.) might bet associated with better training outcomes. There is a clear need to investigate stimulation in conjunction with injury and the role that it may play in network recovery. It is often assumed that concussed patients should limit exposure to any form of stimulation because it mitigates their symptoms; however, targeted stimulation may instead help resolve chronic deficits [79]. Relatedly, the functional connectivity characteristics of our hippocampal network should be examined more completely, as we may discover a structurally modified network achieves nearly the same functional organization that appeared before injury. This analysis would enable us to address how closely functional networks reflect underlying structural connectivity at the microcircuit scale. At the macroscale, a link between axonal tractography and a resting state functional network is established [80,81], but the relationship between structural and functional connectivity is not well understood in microcircuits. Further, characterizing functional networks from these simulations would offer an opportunity to link this work with experimental results measured via microelectrode arrays and make structurally based insights about those empirical functional data [82].

With this work, we investigate the modular network structure of a computational model of the hippocampus, a region of the brain that has well-characterized anatomy and electrophysiology, and we examine the functional implications of plasticity impairment on network-defined pattern separation. These studies contribute to a growing body of work regarding the circuit-level effects of cellular damage in neuronal networks [31,36,37,83]. Studying a posited substrate of physiological learning with this biologically inspired computational model of the hippocampus, which is known for its role in learning and memory, guides new insights into both temporary and more permanent impairments that could occur from cellular-based changes after traumatic injury. In addition, combining this cellular-level mechanistic insight with new tools in data science (e.g., deep learning and machine learning) provides an opportunity to create biologically inspired autonomous learning models that could aid the recovery and repair of damaged circuits. By understanding network-based learning in this hippocampal circuit, we will not only advance practical analytical tools, but we may also develop targeted interventions to improve outcomes for patients with diseases of brain-network organization.

## Acknowledgements

The following figures or figure panels were created with BioRender.com: Fig 1A, Fig 1B, Fig 2, Fig 6A, and Fig 7A.

